# Identification of activity cliffs in structure-activity landscape of androgen receptor binding chemicals

**DOI:** 10.1101/2022.12.06.519328

**Authors:** R.P. Vivek-Ananth, Ajaya Kumar Sahoo, Shanmuga Priya Baskaran, Janani Ravichandran, Areejit Samal

## Abstract

Androgen mimicking environmental chemicals can bind to Androgen receptor (AR) and can cause severe effects on the reproductive health of males. Predicting such endocrine disrupting chemicals (EDCs) in the human exposome is vital for improving current chemical regulations. To this end, QSAR models have been developed to predict androgen binders. However, a continuous structure-activity relationship (SAR) wherein chemicals with similar structure have similar activity does not always hold. Activity landscape analysis can help map the structure-activity landscape and identify unique features such as activity cliffs. Here we performed a systematic investigation of the chemical diversity along with the global and local structure-activity landscape of a curated list of 144 AR binding chemicals. Specifically, we clustered the AR binding chemicals and visualized the associated chemical space. Thereafter, consensus diversity plot was used to assess the global diversity of the chemical space. Subsequently, the structure-activity landscape was investigated using SAS maps which capture the activity difference and structural similarity among the AR binders. This analysis led to a subset of 41 AR binding chemicals forming 86 activity cliffs, of which 14 are activity cliff generators. Additionally, SALI scores were computed for all pairs of AR binding chemicals and the SALI heatmap was also used to evaluate the activity cliffs identified using SAS map. Finally, we provide a classification of the 86 activity cliffs into six categories using structural information of chemicals at different levels. Overall, this investigation reveals the heterogeneous nature of the structure-activity landscape of AR binding chemicals and provides insights which will be crucial in preventing false prediction of chemicals as androgen binders and developing predictive computational toxicity models in the future.

## 1. Introduction

Androgen receptor (AR) is a ligand-dependent nuclear transcription factor mediated by male sex hormones dihydrotestosterone (DHT) and testosterone which are also known as androgens (Davey and Grossmann, 2016; Tan et al., 2015). The binding of these androgens with AR plays a key role in the development of male reproductive system and secondary sexual characteristics (Davey and Grossmann, 2016; Tan et al., 2015). Apart from the androgens, several endocrine disrupting chemicals (EDCs) have been reported to bind with AR and interfere with the normal functioning of the hormones (Jeng, 2014; Rehman et al., 2018; Rodprasert et al., 2021). The modes of action of the EDCs on the human endocrine system are manifold, and these include blocking the binding of the hormones to their native receptors (Karthikeyan et al., 2021, 2019; UNEP, 2013). These EDCs have a deleterious effect on human reproductive health which includes developmental abnormalities in the reproductive tract, poor semen quality and testicular cancer (Rehman et al., 2018; Rodprasert et al., 2021). Among the myriad environmental chemicals in the human exposome, it is therefore imperative to identify the EDCs which can bind with AR and interfere with the normal functioning of the male hormones.

Structure-activity relationship (SAR) based analysis can help in predicting potential AR binding chemicals. Such approaches have been undertaken previously to explore the SAR of AR binding chemicals (Fang et al., 2003). Yet, to develop highly predictive quantitative SAR (QSAR) models from the SAR data, it is important to study the activity landscape of the AR binding chemicals. To the best of our knowledge, no such activity landscape analysis has been performed for the AR binding chemicals to date. Activity landscape analysis of the AR binders can help characterize the activity landscape, in particular, the identification of continuous and discontinuous SARs including the activity cliffs (Bajorath, 2017; Cruz-Monteagudo et al., 2014; Maggiora, 2006). Activity cliffs are a unique feature of an activity landscape, wherein the SAR becomes discontinuous, as these activity cliffs are chemicals which are structurally highly similar but with high activity difference (Bajorath, 2017; Cruz-Monteagudo et al., 2014; Maggiora, 2006). Notably, the activity cliffs should not be considered as outliers in the SAR data as they are inherent features of the structure-activity landscape being analyzed (Cruz-Monteagudo et al., 2014; Maggiora, 2006). Thus, the activity landscape analysis and identification of the activity cliffs in the space of AR binding chemicals will enable the creation of better predictive models for EDCs.

Multiple computational methods for performing activity landscape analysis and identification of activity cliffs have been developed in the cheminformatics literature including by Medina-Franco and colleagues, Guha and Van Drie, and Bajorath and colleagues (Guha and Van Drie, 2008; Hu and Bajorath, 2012; Medina-Franco et al., 2009; Méndez-Lucio et al., 2012; Naveja et al., 2018; Naveja and Medina-Franco, 2015; Peltason and Bajorath, 2007; Wawer et al., 2008). Medina-Franco and colleagues have extensively used the two-dimensional (2D) representation of activity landscape, Structure-Activity Similarity (SAS) map, which considers the structural similarity and activity difference of chemicals to identify the activity cliffs (Méndez-Lucio et al., 2012; Naveja et al., 2018; Naveja and Medina-Franco, 2015). SAS map based approach to computationally detect activity cliffs was first proposed by Shanmugasundaram and Maggiora in 2001, and has since been used to study both global and local landscape for large chemical datasets (Naveja et al., 2018; Naveja and Medina-Franco, 2015; Shanmugasundaram and Maggiora, 2001). Further, Guha et al. have developed ‘SALI’ scoring based approach to numerically quantify activity cliffs in a given dataset (Guha and Van Drie, 2008). Though these computational approaches to analyze the activity landscape have been extensively used on chemical datasets for drug discovery research, the use of these methods to find activity cliffs in datasets of toxic chemicals is limited. A notable exception is the work of Naveja et al. wherein SAS map approach was used to computationally detect activity cliffs and ‘activity cliff generators’ in a dataset of 121 estrogen receptor binding chemicals (Naveja et al., 2018). Further, Naveja et al. provided a mechanistic interpretation of the difference in the binding affinities of the activity cliffs (Naveja et al., 2018).

In this study, we performed a systematic analysis of the activity landscape of a curated list of 144 AR binding chemicals with reported AR binding affinity from Fang et al. to identify the activity cliffs and activity cliff generators (Fang et al., 2003). We performed chemical space exploration, clustering and diversity analysis of the curated dataset of 144 AR binding chemicals using multiple computational approaches. Further, we studied both the global and local landscape of the curated dataset using SAS map based approach. Moreover, we computed the SALI score for all pairs of chemicals and visualized it using SALI heatmap. By overlaying the activity cliffs identified from the SAS map on the SALI heatmap, we also compared the two methodologies for identifying activity cliffs. Lastly, using the structural information of chemicals at different levels, we also provide a systematic structural classification of the activity cliff pairs identified in this study.

## 2. Methods

### 2.1. Chemical dataset curation and annotation

Previously, Fang et al. had experimentally determined the androgen receptor (AR) binding affinity of 202 natural, synthetic and environmental chemicals against recombinant rat AR protein using competitive receptor binding assay (Fang et al., 2003). In particular, the Fang et al. dataset provides the CAS identifiers, experimentally determined half maximal inhibitory concentration (IC_50_), and the relative binding affinity (RBA) for the 202 chemicals. Moreover, Fang et al. classified the chemicals into 14 broad classes namely, steroids, diethylstilbestrols (DESs), phytoestrogens, phenols, flutamides, diphenylmethanes, polychlorinated biphenyls (PCBs), organochlorines, phthalates, aromatic hydrocarbons, noncyclic chemicals, aromatic acids, phenol like chemicals, and others.

Here, we leveraged the Fang et al. dataset to study the SAR landscape of AR binding chemicals. First, we removed chemicals which were labeled as “nonbinders” and “slight binders” in Fang et al. (Fang et al., 2003). Afterwards, we collected the two-dimensional (2D) chemical structures for the remaining 146 AR binders from ChemIDplus (https://chem.nlm.nih.gov/chemidplus/). The compiled chemical structures were processed using a cleaning protocol which included removing the invalid structures, duplicate structures and salts using MayaChemTools (Sud, 2016). Further, we removed two acyclic chemicals from the dataset. In consequence, we curated a dataset of 144 natural, synthetic and environmental chemicals (Supplementary Table S1), along with their AR binding affinities and chemical structures starting from the information in Fang et al. (Fang et al., 2003). Lastly, using the ClassyFire webserver, we structurally classified the 144 chemicals (Supplementary Table S1) in our dataset (Djoumbou Feunang et al., 2016).

### 2.2. Chemical structure characterization

We characterized the structures of the 144 chemicals in our dataset using structural fingerprints, physicochemical properties and molecular scaffolds. First, we used ECFP4 fingerprints implemented in RDKit (Morgan, 1965; Rogers and Hahn, 2010) to capture the structural features of the 144 chemicals. Thereafter, we used ECFP4 fingerprints to compute pairwise chemical structure similarity for the 144 chemicals. Second, we computed six physicochemical properties (PCP) namely, hydrogen bond donors (HBD), hydrogen bond acceptors (HBA), octanol/water partition coefficient (LogP), molecular weight (MW), topological polar surface area (TPSA) and number of rotatable bonds (RTB) for the 144 chemicals. Third, we computed the molecular scaffolds for the 144 chemicals using the Bemis-Murcko definition (Supplementary Table S1) (Bemis and Murcko, 1996).

### 2.3. Structure based clustering

We analyzed the global and local structure-activity relationship (SAR) for the 144 chemicals by clustering the chemicals based on structural similarity. To cluster the chemicals based on structural similarity, we first computed the pairwise chemical structure similarity for all pairs of chemicals in the library. The chemical structure similarity between any two chemicals was computed via Tanimoto coefficient (Tc) (Tanimoto, T. T., 1957) using ECFP4 fingerprints. Thereafter, we constructed a similarity matrix for the whole library using the Tc values for all pairs of chemicals in our dataset.

In order to visualize the high-dimensional dataset, we used principal component analysis (PCA) (Jolliffe, 1986) to project the data to two dimensions. Notably, the first two principal components (PC1; PC2) capture 53.15% variance in the whole library. From the PCA plot (Figure 1), we find that the 144 chemicals in the whole library can be grouped into three clearly separated clusters.

**Figure 1:**
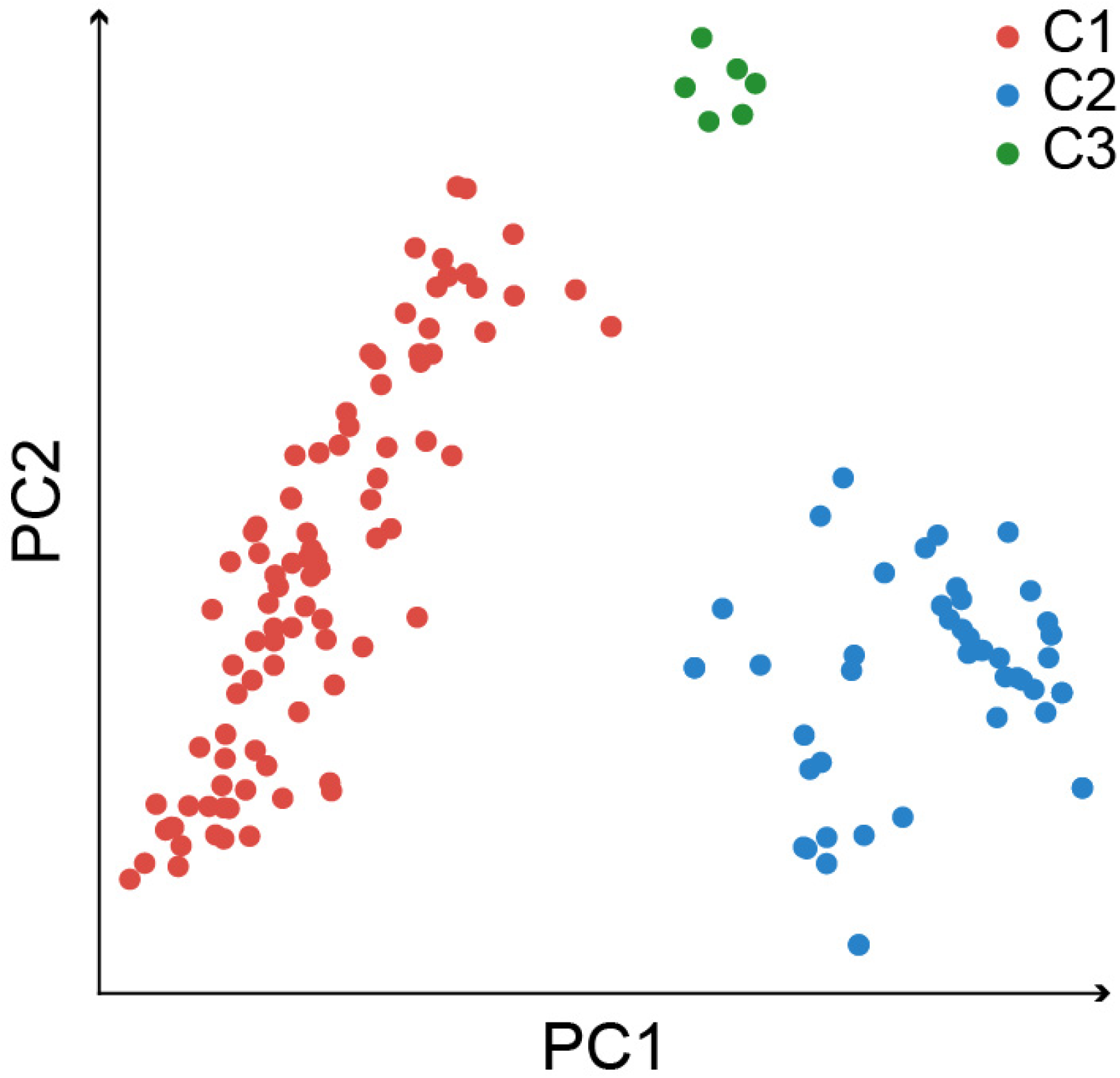
Principal Component Analysis (PCA) of the 144 AR binding chemicals. PCA was performed on the similarity matrix which encapsulates the pairwise structural similarities between all pairs of chemicals. The data points in the two-dimensional PCA plot (PC1; PC2) corresponds to the 144 chemicals in the whole library, and the data points are colored based on the three chemical clusters identified from the chemical similarity network (CSN).

To further assist the clustering of chemicals in our dataset, we constructed a chemical similarity network (CSN) of the 144 chemicals. The nodes in the CSN represent the chemicals and the edges are drawn between two nodes if the Tc between the respective chemical pair is ≥ 0.2 (Figure 2). The CSN was visualized using Gephi software package version 0.9.7 (Bastian et al., 2009). Thereafter, we used the Louvain community detection (with resolution parameter set to 5.0) within Gephi package to identify clusters of chemicals within the CSN (Blondel et al., 2008). Lastly, we colored the data points in the PCA plot (Figure 1) based on the clusters identified within the CSN (Figure 2; Supplementary Table S1).

**Figure 2:**
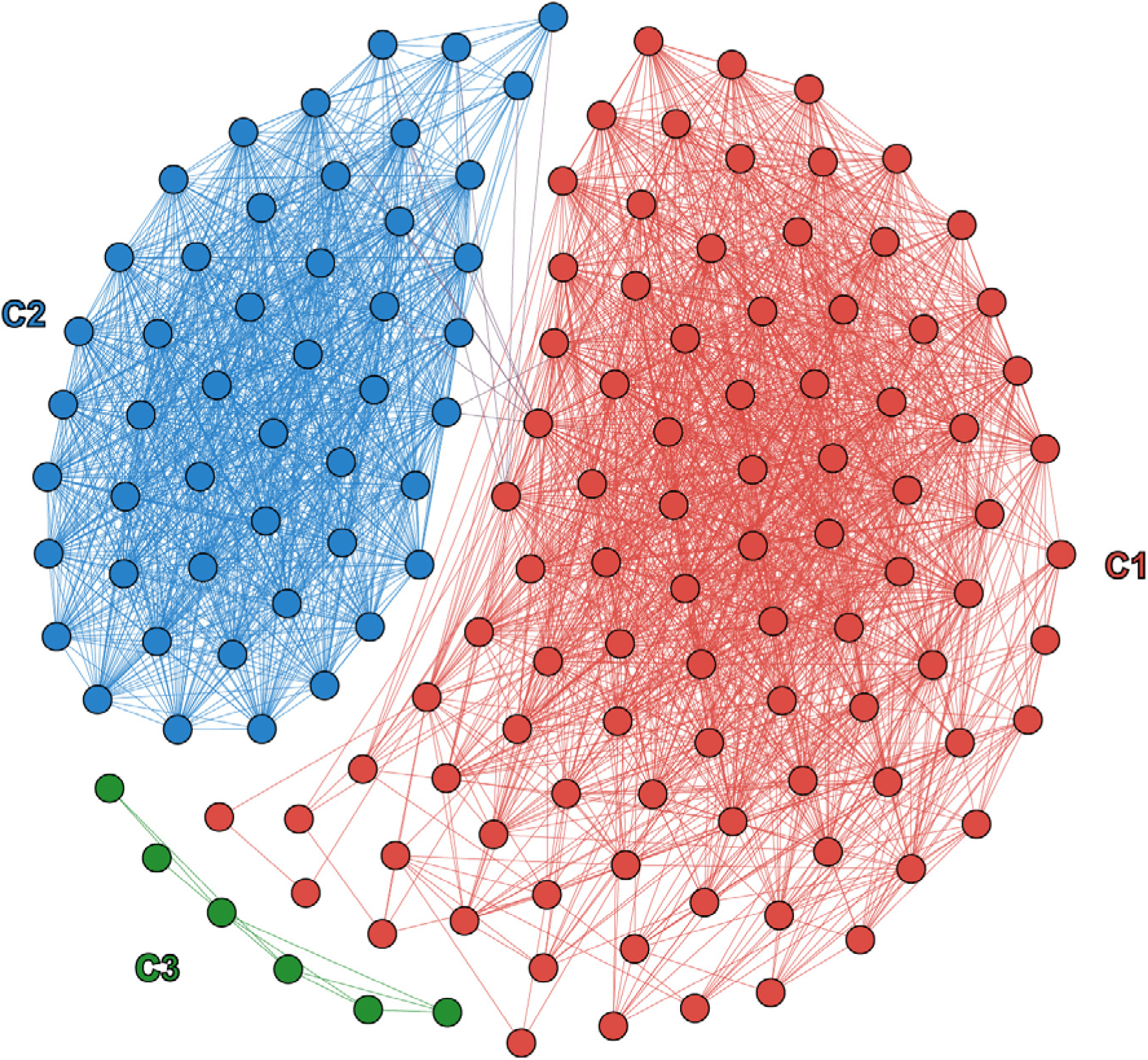
Chemical Similarity Network (CSN) of the 144 AR binding chemicals. The CSN was constructed based on the pairwise structural similarity computed via Tc using ECFP4 fingerprints. Edges were assigned between two nodes if Tc ≥ 0.2. The nodes and edges belonging to the chemical clusters C1, C2 and C3 are colored in red, blue and green, respectively. The edges between clusters C1 and C2 are colored in shades of magenta.

### 2.4. Global diversity

The Consensus Diversity Plot (CDP) (González-Medina et al., 2016; Naveja et al., 2018) helps in visualizing and comparing the diversity of different chemical libraries. We used CDP to compare the diversity of the three chemial clusters and the whole library of 144 chemicals analyzed here (Figure 4). The x-axis of the CDP represents median Tc obtained using MACCS keys fingerprints, capturing the structural diversity. The y-axis of the CDP represents the area under the curve (AUC) obtained from the cyclic system retrieval curve (CSR) (Lipkus et al., 2008; Vivek-Ananth et al., 2022), capturing the scaffold diversity. The color of the data points in the CDP represents the PCP diversity of the clusters or the whole library. Note, red color indicates high PCP diversity and blue color indicates low PCP diversity (González-Medina et al., 2017; Vivek-Ananth et al., 2022). PCP diversity is the mean Euclidean distance between all pairs of chemicals in a cluster or the whole library computed using the six physicochemical properties. Finally, the size of a data point in the CDP represents the relative size of a cluster or the whole library.

**Figure 3:**
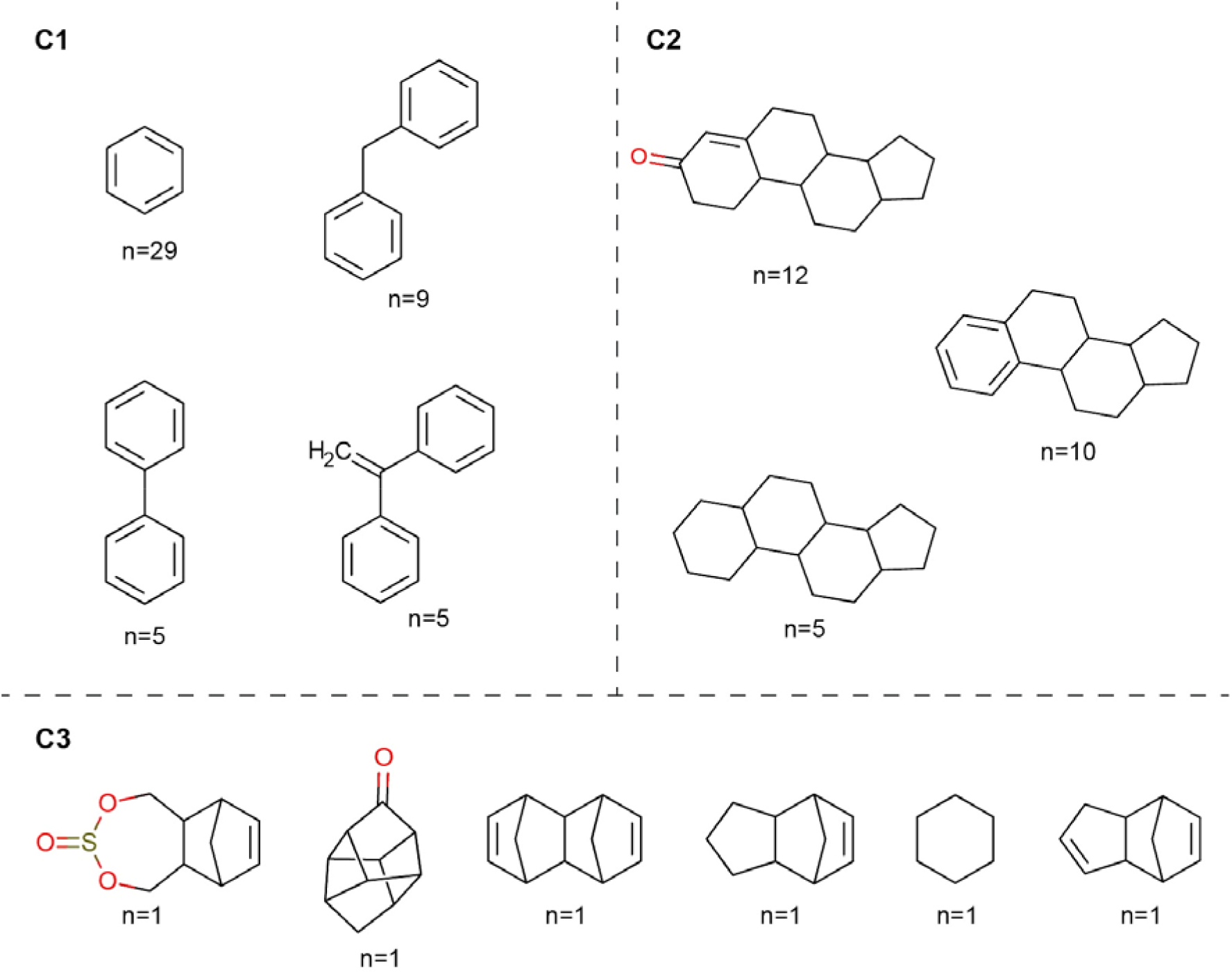
Top three Bemis-Murcko scaffolds for the three chemical clusters of AR binding chemicals in terms of the frequency (*n*) of their occurrence in each chemical cluster.

**Figure 4:**
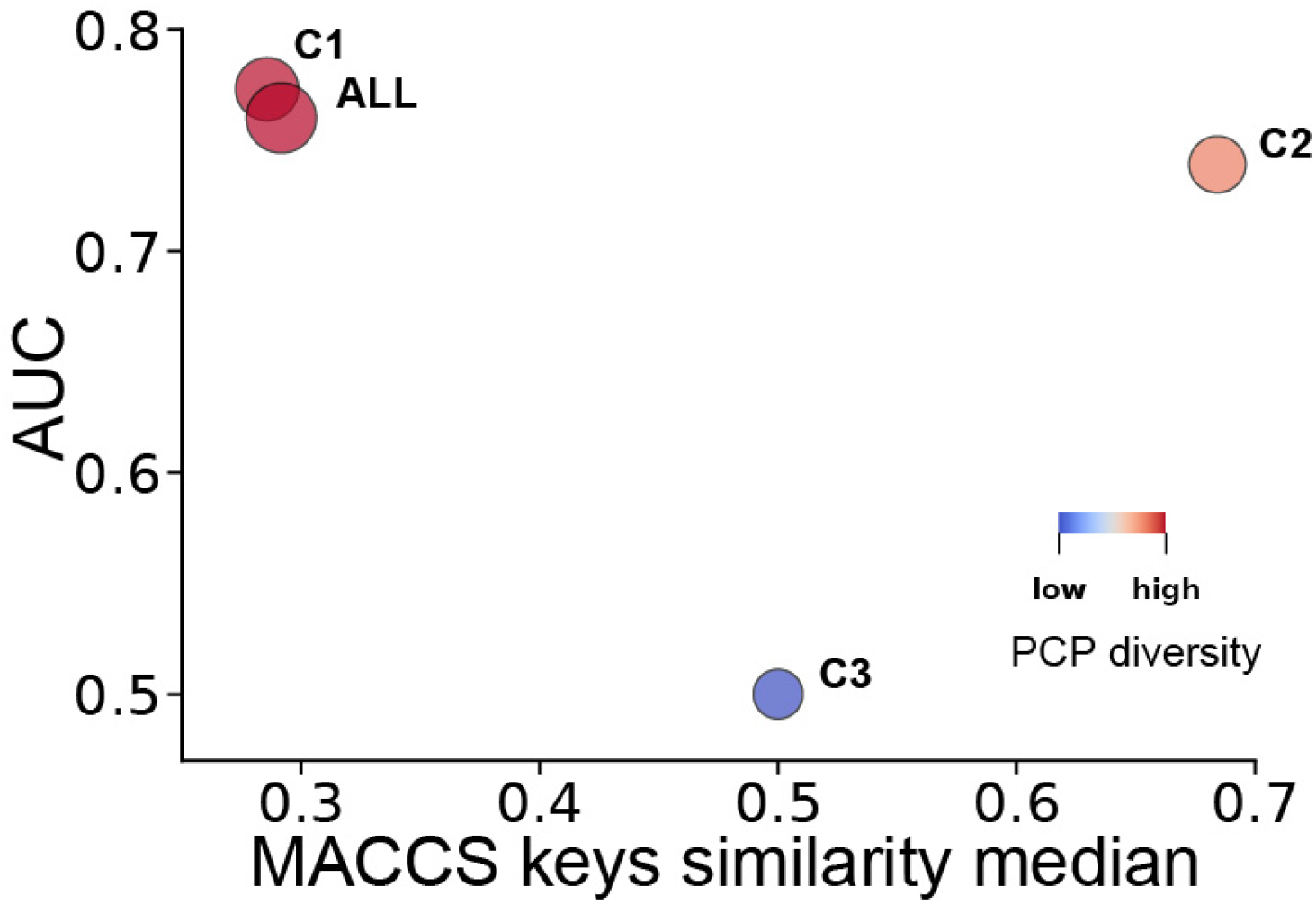
Consensus Diversity Plot (CDP) depicting the global diversity of the three chemical clusters C1, C2, and C3, and the whole library, ALL. The x-axis of the CDP represents the median Tc obtained using MACCS keys fingerprints and the y-axis represents AUC. The PCP diversity computed from the mean Euclidean distance using the six physicochemical properties are represented by the color of the data points: red represents high PCP diversity whereas blue represents low PCP diversity. The relative size of the dataset is represented by the size of the data points.

### 2.5. Activity difference

For any pairs of chemicals in a cluster or the whole library, the absolute value of pairwise activity difference was calculated using the formula:

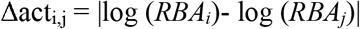

where, *RBA_i_* and *RBA_j_* are the relative binding affinities of the *i*^th^ and *j*^th^ chemicals, respectively as determined by Fang et al. (Fang et al., 2003).

### 2.6. Activity cliff identification

Activity landscape analysis helps in studying the nature of the SAR of a chemical space. In particular, ‘activity cliffs’ or discontinuous SAR identified by the analysis can help in capturing key pharmacophore regions necessary for biological activity. Activity landscape analysis can be done using various approaches. Here, we used Structure-Activity Similarity (SAS) map (Naveja et al., 2018; Shanmugasundaram and Maggiora, 2001) and Structure-Activity Landscape Index (SALI) (Guha, 2012; Guha and Van Drie, 2008) for the identification of activity cliffs.

#### 2.6.1. Structure-Activity Similarity (SAS) map

Here, we generated SAS maps for the three clusters and the whole library of 144 chemicals (Figure 5). Briefly, the SAS map has 4 quadrants (I to IV). Quadrant I denotes scaffold hopping region (with chemicals having low structural similarity and low activity difference), quadrant II corresponds to smooth region of the SAR space (with chemicals having high structural similarity and low activity difference), quadrant III identifies activity cliffs (with chemicals having high structural similarity and high activity difference), and quadrant IV represents an uncertain region (with chemicals having low structural similarity and high activity difference). Importantly, quadrant III of the SAS map is the region of interest in this work.

**Figure 5:**
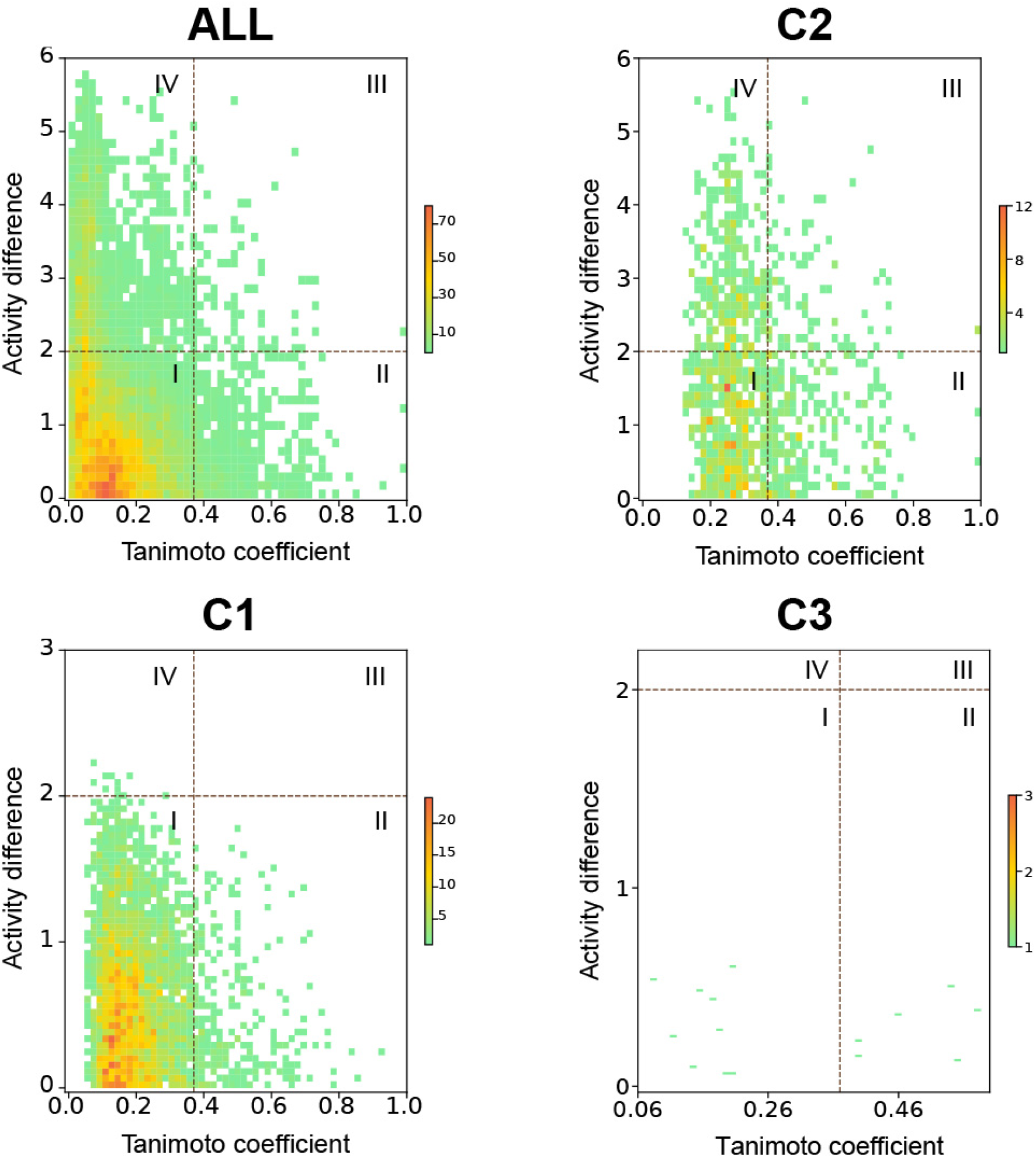
Structure-Activity Similarity (SAS) map for the whole library (ALL) and three chemical clusters (C1, C2 and C3). In each case, the SAS map is divided into 4 quadrants (denoted by I, II, III and IV) by considering x-axis threshold as 0.37 and y-axis threshold as 2 logarithmic units (Methods). Regions in each plot are colored based on the number of data points in that region (green for the low dense region and red for the high dense region).

The x-axis of the SAS map represents the Tc obtained using ECFP4 fingerprints for different pairs of chemicals. The y-axis of the SAS map represents the absolute activity difference between different pairs of chemicals (Δact_i,j_). The median plus two standard deviations for the distribution of Tc obtained using ECFP4 fingerprints was computed for all pairs of chemicals in the whole library, and thereafter, it was used as x-axis threshold to demarcate the quadrants in the SAS map. The x-axis threshold for the whole library of 144 chemicals was determined to be 0.37. Note that the same x-axis threshold was used for both global and local SAS maps. Following Naveja et al., the y-axis threshold was set at 2 logarithmic units or 100-fold change in the activity difference (Naveja et al., 2018). We remark that SAS maps are two-dimensional heatmaps wherein a continuous color gradient is used to represent the number of data points in a region.

#### 2.6.2. Structure-Activity Landscape Index (SALI)

SALI can be used to quantitatively characterize activity landscapes (Guha and Van Drie, 2008). We computed the SALI scores for all pairs of chemicals in a cluster and the whole library. SALI score is based on the activity difference and pairwise structural similarity, and is calculated as follows:

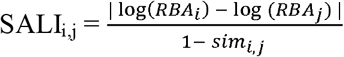

where *RBA_i_* and *RBA_j_* are the relative binding affinities of the *i*^th^ and *j*^th^ chemicals and *sim*_i,j_ is the pairwise structural similarity between *i*^th^ and *j*^th^ chemicals. Note, when the *sim*_i,j_ value for a pair of chemicals becomes 1, the SALI score becomes undefined. For such a pair of chemicals, the SALI score is assigned equal to that of the chemical pair with the maximum SALI score in the dataset. Tc using ECFP4 fingerprints was used for computing pairwise structural similarity of chemicals. Pairs of chemicals with high values of SALI correspond to activity cliffs and the computed scores were visualized using the SALI heatmap (Guha and Van Drie, 2008). Importantly, we highlight the activity cliffs identified from the SAS map by marking them as black boxes in the SALI heatmap (Figure 6). Notably, this combined approach helps in visually identifying the overlap between the activity cliffs from SAS map and chemical pairs with high SALI scores. The plots, including heatmaps were generated via in-house python scripts using Matplotlib package (Hunter, 2007).

**Figure 6:**
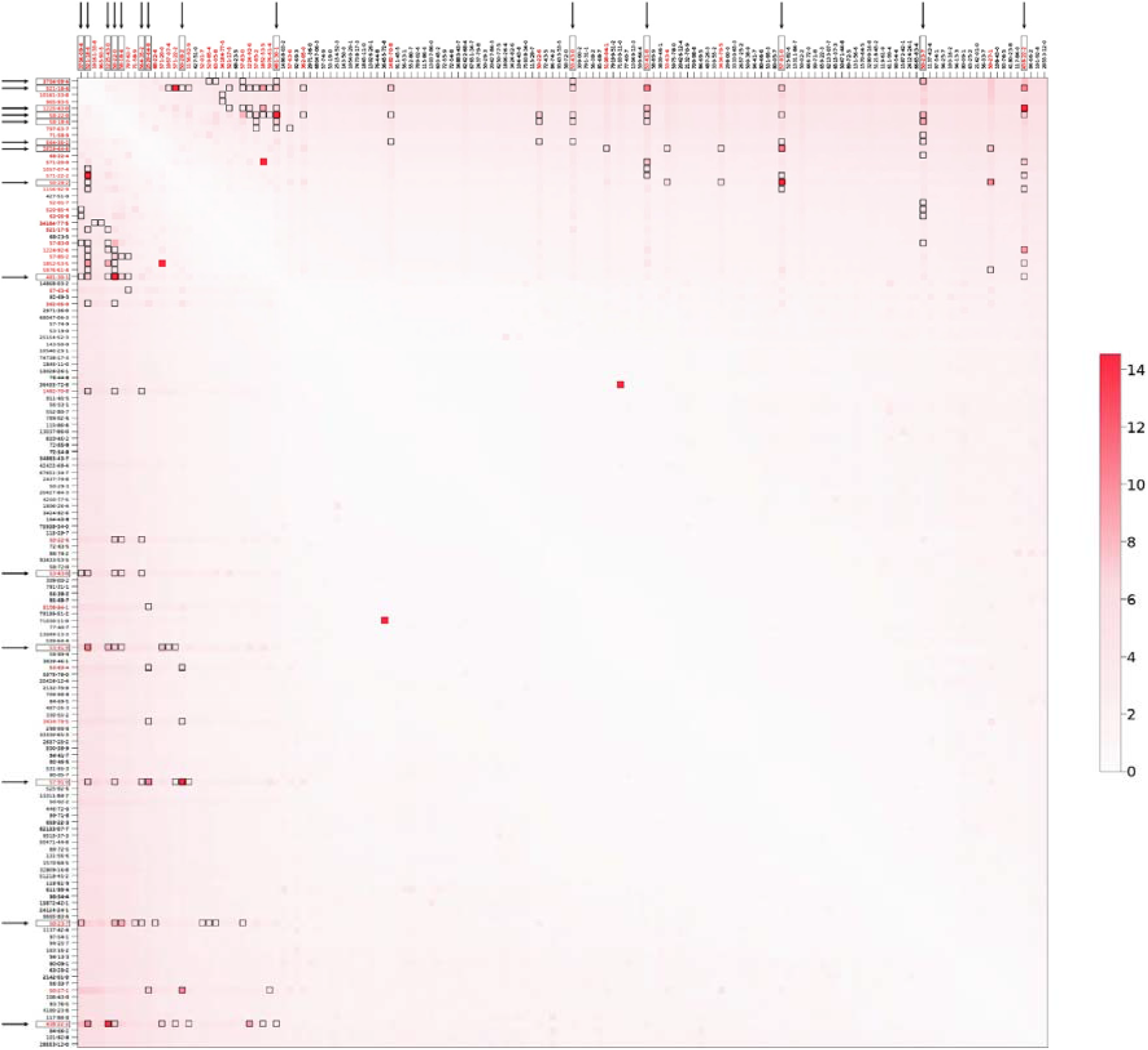
SALI score based heatmap for all chemical pairs of the 144 AR binding chemicals. The x-axis and y-axis of the heatmap are labelled with CAS identifiers of the 144 chemicals arranged from left to right and top to bottom in the descending order of their respective AR binding affinities. Each cell in the heatmap represents the chemical pair and is colored based on the computed SALI score: darker the cell, higher the SALI score. The chemical pairs (cells in heatmap) identified as activity cliffs in the SAS map are highlighted with black boxes. The CAS identifiers of the 41 chemicals which makeup the 86 activity cliffs in the SAS map are shown in red color. Further, the labels of the 14 ACGs identified in the SAS map are boxed and marked with arrows.

## 3. Results

### 3.1. Visualization and exploration of the chemical space of AR binders

Figure 1 displays the two-dimensional PCA plot generated from the similarity matrix for the whole library of 144 AR binding chemicals (Methods). In the PCA plot, we observed three chemical clusters, spatially separated in the two-dimensional space obtained using the first two principal components (PC1; PC2) (Figure 1). Further, we built the Chemical Similarity Network (CSN) of the 144 AR binding chemicals (Methods). Applying Louvain community detection method on the CSN, we again identified three chemical clusters (Supplementary Table S1) which are more likely to contain structurally similar chemicals (Figure 2) (Djoumbou Feunang et al., 2016). We annotated the data points in the PCA plot using three different colors corresponding to the three chemical clusters identified in the CSN (Figures 1 and 2). From Figure 1, we can see that the cluster information obtained from the CSN concurs with the clustering obtained from the PCA plot. The chemical cluster C1 contains 90 chemicals (62.5% of the whole library), C2 contains 48 chemicals (33.33% of the whole library) and C3 contains 6 chemicals (4.17% of the whole library).

Moreover, we annotated the chemicals in the three clusters using the chemical class information predicted from ClassyFire (Supplementary Table S1), and their occurrence in the list of endocrine disrupting chemicals (EDCs) compiled in DEDuCT and publicly available lists on chemical regulation (Djoumbou Feunang et al., 2016; Karthikeyan et al., 2021, 2019). Firstly, we find that the 144 chemicals in the whole library belong to 21 chemical classes. The chemicals in cluster C1 belong to a diverse set of chemical classes. ‘Benzene and substituted derivatives’ (47 of 90 chemicals) is the most prevalent chemical class in C1. The chemicals in cluster C2 are dominated by the chemical class ‘Steroids and steroid derivatives’ (44 of 48 chemicals). The chemicals in cluster C3 belong to diverse chemical classes. Importantly, we observed that the chemical classes of each cluster are unique, i.e. there is no overlap among the clusters in terms of the occurrence of chemical classes (Supplementary Table S1).

Secondly, we find 62 chemicals in the whole library are EDCs with documented adverse health effects based on a comparative analysis with the list of 792 potential EDCs compiled in DEDuCT (Karthikeyan et al., 2021). The distribution of these 62 EDCs among the chemical clusters C1, C2 and C3 was found to be 46, 11 and 5 chemicals, respectively. Thirdly, to comprehend the regulatory status of the AR binding chemicals in the whole library, in view of their potential adverse health effects on humans, we considered six publicly available lists on chemical regulations. We find that, among the 144 AR binding chemicals, 31 are present in ‘California Proposition 65 (CP65)’ (https://oehha.ca.gov/proposition-65/proposition-65-list) list, 7 are present in ‘Restricted substances under REACH’ (https://echa.europa.eu/substances-restricted-under-reach) list, 8 are present in ‘SVHC under REACH’ (https://echa.europa.eu/substances-of-very-high-concern-identification-explained) list, 2 are present in ‘Toxic chemicals restricted to be imported or exported in China’ (http://www.cirs-reach.com/China_Chemical_Regulation/Registration_of_import_export_of_toxic_chemicals_in_China.html) list, 18 are present in ‘EU list of substances prohibited in cosmetic products’ (https://eur-lex.europa.eu/legal-content/EN/TXT/?uri=celex:02009R1223-20150416) list, and 10 are present in ‘Schedule 1 hazardous chemicals list in India’ (http://moef.gov.in/wp-content/uploads/2019/08/SCHEDULE-I.html) list. In total, 42 of the 144 AR binding chemicals are present in at least one of the six chemical regulations considered here. Further, of the 62 chemicals in the whole library identified as potential EDCs, 29 are present in at least one of the six chemical regulations considered here. We remark that this dataset of 144 AR binding chemicals containing potential EDCs and regulated chemicals of concern can serve as a benchmark dataset to study the local and global SAR of the AR binding chemicals.

### 3.2. Scaffold content of the AR binding chemicals

In order to explore the scaffold content, we computed the Bemis-Murcko scaffolds for each of the 144 AR binding chemicals (Methods). Figure 3 shows the top 3 scaffolds in terms of the frequency of its occurrence in each chemical cluster. The number of unique molecular scaffolds in the chemical clusters C1, C2 and C3 are 29, 21 and 6, respectively. The scaffold content of cluster C1 is dominated by the benzene scaffold (i.e., present in 29 chemicals). The scaffold content of cluster C2 is dominated by scaffolds belonging to the steroid class. In cluster C3, all the 6 chemicals have a unique scaffold. Of note, the scaffold content of each cluster is unique, i.e. there is no overlap among the clusters in terms of the scaffolds of the chemicals (Supplementary Table S1). The presence of non-overlapping molecular scaffolds and chemical classes among the three clusters further justifies the clustering of the chemical space of the 144 AR binders into three clusters. Therefore, we used these chemical clusters and the whole library to study the local and global SAR of the AR binding chemicals.

### 3.3. Consensus Diversity Plot (CDP) and global diversity of AR binding chemicals

Figure 4 shows the CDP which helps in analyzing the global diversity of the chemical clusters and the whole library of AR binders (Methods). The data points in CDP namely, C1, C2, C3 and ALL correspond to the three chemical clusters and the whole library, respectively. The data points in the left of the CDP have high structural diversity, those in the bottom of the CDP have high scaffold diversity, and data points colored in red have high PCP diversity. From the CDP, we observed that the chemicals in cluster C1 have high structural diversity based on the median of Tc obtained using the MACCS keys fingerprints and low scaffold diversity based on AUC (Figure 4). We can see from Figure 4 that the whole library (ALL) has high structural diversity and low scaffold diversity. The chemical cluster C1 also has similar global diversity to the ALL, possibly since C1 accounts for 62.5% of the chemicals in the whole library (Table 1). The low structural diversity of C2 suggests that the chemicals in C2 are structurally very similar. The chemical cluster C2 has AUC similar to C1 and ALL, suggesting that C1, C2 and ALL have similar scaffold diversity (Figure 4; Table 1). The chemical cluster C3 has the highest scaffold diversity with AUC value of 0.5. This is because all the six chemicals in C3 have unique molecular scaffolds. Relatively speaking, C1 has highest PCP diversity and C3 has the lowest PCP diversity (Figure 4; Table 1).

**Table 1:** Global diversity of the three chemical clusters, C1, C2, and C3, and the whole library, ALL. For each set, the table lists the number of chemicals present, median Tc using MACCS keys fingerprints, AUC and the PCP diversity.

| Chemical dataset | Number of chemicals | Median Tc using MACCS keys fingerprints | AUC | PCP diversity |
| --- | --- | --- | --- | --- |
| ALL | 144 | 0.29 | 0.76 | 3.14 |
| C1 | 90 | 0.29 | 0.77 | 3.13 |
| C2 | 48 | 0.68 | 0.74 | 2.95 |
| C3 | 6 | 0.5 | 0.5 | 2.13 |

### 3.4. Activity landscape analysis to explore the SAR of the AR binding chemicals

Activity landscape analysis is widely used to analyze the structure-activity relationship (SAR) of diverse chemical spaces (Wassermann et al., 2010). Structure-Activity Similarity (SAS) map is the first method to be developed for 2D visual representation of activity landscape (Shanmugasundaram and Maggiora, 2001). SAS map takes into account the structural similarity and activity difference of all possible pairs of chemicals in a given dataset. A SAS map is a 2D heatmap with continuous color gradient reflecting the number of data points in a region (Methods). In particular, the color gradient ranges from green for the low density regions to red for high density regions (Figure 5). Figure 5 shows the SAS maps for the whole library (ALL), and the three chemical clusters (C1, C2, and C3). For the whole library (ALL), the majority of the chemical pairs are in region I (Figure 5; Table 2). This suggests that the corresponding chemical pairs are structurally diverse with low activity difference. Notably, we find only 86 pairs of chemicals in region III. Region III contains pairs of chemicals with high structural similarity and high activity difference, i.e., the activity cliffs. In particular, the 86 activity cliff pairs (Supplementary Table S2) in region III are formed by 41 chemicals.

**Table 2:**
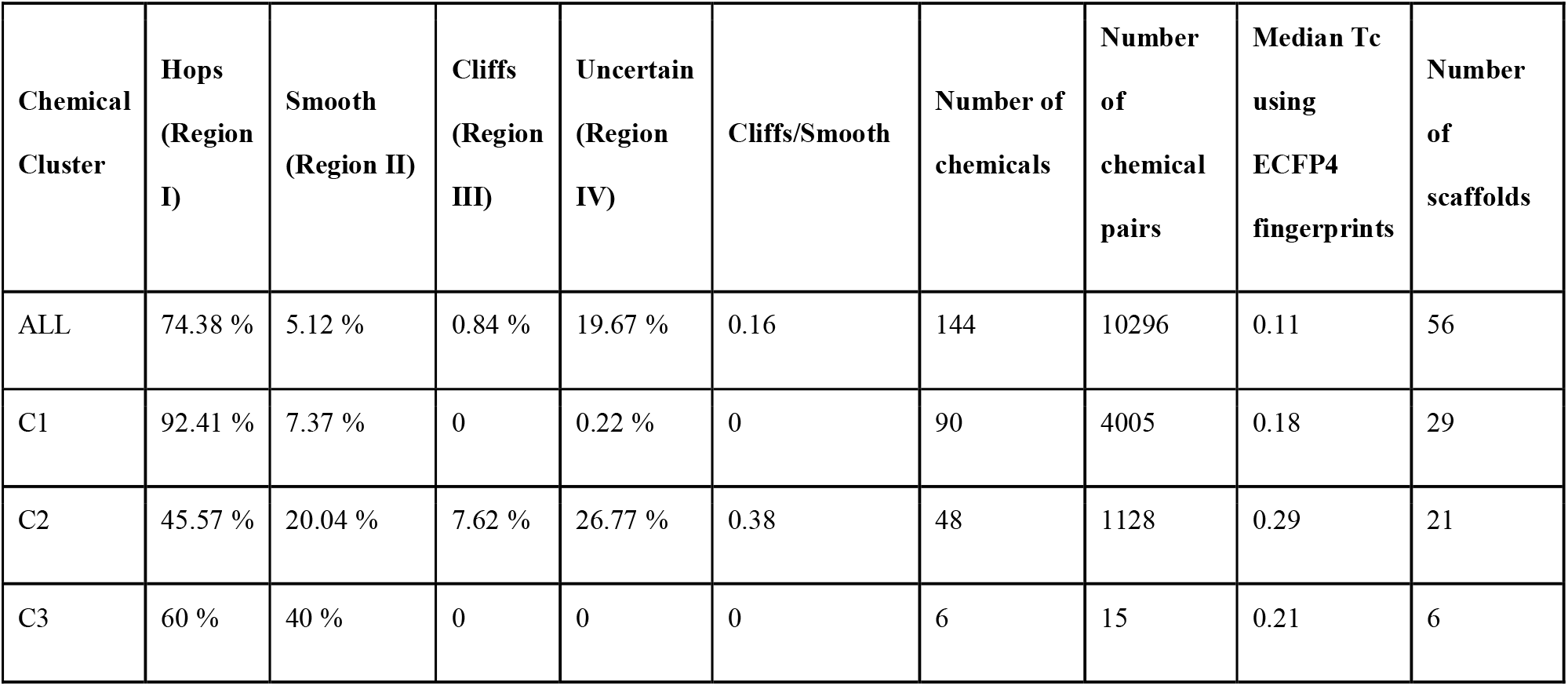
The table summarises the percentage of chemical pairs in the four regions or quadrants of the SAS maps for the whole library (ALL) and the three chemical clusters (C1, C2 and C3). The table also provides the ratio of the chemical pairs present in the activity cliff region (quadrant III) to the chemical pairs present in the smooth region (quadrant II). Further, the table provides the number of chemicals present in the whole library and three chemical clusters, the number of chemical pairs, median Tc computed using the ECFP4 fingerprints, and the number of unique molecular scaffolds in the whole library and the three chemical clusters.

Upon analyzing the local SAS maps for the three clusters, we found that all of these 86 activity cliff pairs belong to cluster C2. Table 2 provides a quantitative summary of the percentage of chemical pairs in the different regions of the SAS map for the three chemical clusters and the whole library. As noted above, the cluster C2 is the only cluster with activity cliffs with 7.62% of the chemical pairs in activity cliff region. In other words, the two other clusters C1 and C3 do not have activity cliff pairs. The local activity landscape for cluster C2 is rough and heterogeneous since C2 has chemical pairs in both smooth and activity cliff regions (Methods). We find that the 41 AR binding chemicals that form activity cliff pairs in cluster C2 belong to the chemical class ‘Steroids and steroid derivatives’.

Further, we identified the activity cliff generators (ACGs) among the 86 activity cliff pairs using the criterion proposed by Naveja et al. (Naveja et al., 2018). We considered a chemical to be an ACG if it is part of at least 5 activity cliff pairs. Following this criterion, we identified 14 chemicals as ACGs (Supplementary Table S3). Notably, we find that the two principal androgens, DHT and testosterone form the maximum number of activity cliff pairs (i.e., 17 and 13, respectively). In a later subsection, we provide a detailed classification of the identified activity cliff pairs.

### 3.5. SALI based exploration of the AR binding chemicals

To date, multiple numerical approaches have been proposed to quantify SAR discontinuity or activity cliffs (Cruz-Monteagudo et al., 2014). Structure-Activity Landscape Index (SALI) introduced by Guha et al. is another widely-used method for identifying and quantifying the activity cliffs in a SAR landscape (Guha and Van Drie, 2008). Herein, we followed Guha et al. to compute the SALI scores for all pairs between the 144 AR binding chemicals (Methods). The computed SALI scores for all chemical pairs can be visualized using the SALI heatmap (Guha and Van Drie, 2008). Figure 6 shows the SALI heatmap for all pairs between the 144 AR binding chemicals. Briefly, both x- and y-axis of this heatmap are annotated with the CAS identifiers of the AR binders, and the chemicals are arranged from left to right and top to bottom according to descending order of their AR binding affinities (Figure 6). Further, each cell in the SALI heatmap represents a chemical pair and the cells are colored based on the SALI score of the corresponding chemical pair (darker the cell color, higher the SALI score). Certain cells of the SALI heatmap are marked with black boxes, if the corresponding chemical pairs were identified as activity cliffs from the analysis of the SAS map (Figures 5 and 6; Supplementary Table S2). The CAS identifiers (axes labels) corresponding to the 41 chemicals which makeup the 86 activity cliff pairs (Supplementary Table S2) identified from the SAS map are colored in red. Also, the CAS labels of the 14 ACGs (Supplementary Table S3) identified from the SAS map are boxed and marked with arrows (Figure 6).

From Figure 6, we can see that most of the chemical pairs identified as activity cliffs in the SAS map, also have high SALI scores. Further, we also noted that some chemical pairs having high SALI score (dark colored cell) were not identified as activity cliffs in the SAS map (and thus, not highlighted with black boxes in the SALI heatmap). We find such pairs to be stereoisomers with low activity difference. We note that the chemical fingerprints used to compute structural similarity does not capture the stereoisomer information, and this renders the structural dissimilarity (i.e., denominator in the SALI score definition) to be zero. Due to this, in spite of the low activity difference, the stereoisomer pairs were assigned the highest SALI score (Methods). However, in the case of SAS map, such stereoisomer pairs are located in region II (as they have high similarity and low activity difference), and hence are not identified as activity cliffs. Lastly, we observed that the SALI heatmap cells with intermediate SALI score have not been identified as activity cliffs in the SAS map approach, possibly due to the x- and y-axis thresholds used to demarcate the four regions in the SAS maps (Figure 6). In sum, there is a large overlap between the activity cliff pairs identified from the SAS map and the chemical pairs with high SALI score. These observations further support the 86 activity cliff pairs identified in this study.

### 3.6. Structural classification of Activity cliffs

Previously, Hu et al. have proposed a conceptually different methodology from SAS map or SALI score to identify the activity cliffs (Hu and Bajorath, 2012). The proposed methodology includes the structural classification of chemical pairs with high activity difference, using the molecular scaffolds, R-groups, and topology of the R-groups in the chemical structures. Here, we followed Hu et al. to systematically classify the 86 activity cliff pairs identified using the SAS map.

Figure 7 shows the computational workflow used to structurally classify the activity cliff pairs by considering the structural information of chemicals at different levels. Here we classified the activity cliff pairs into 6 structural categories namely:

i. Chirality cliff: An activity cliff pair having same molecular scaffold, R-groups and topology of R-groups.
ii. Topology cliff: An activity cliff pair having same scaffold and R-groups but different topology of R-groups.
iii. R-group cliff: An activity cliff pair having same scaffold but different R-groups.
iv. Scaffold cliff: An activity cliff pair having different scaffolds, but same R-groups and topology of the R-groups.
v. Scaffold/Topology cliff: An activity cliff pair having different scaffolds and topology of the R-groups, but same R-groups.
vi. Scaffold/R-group cliff: An activity cliff pair having different scaffolds and R-groups.

**Figure 7:**
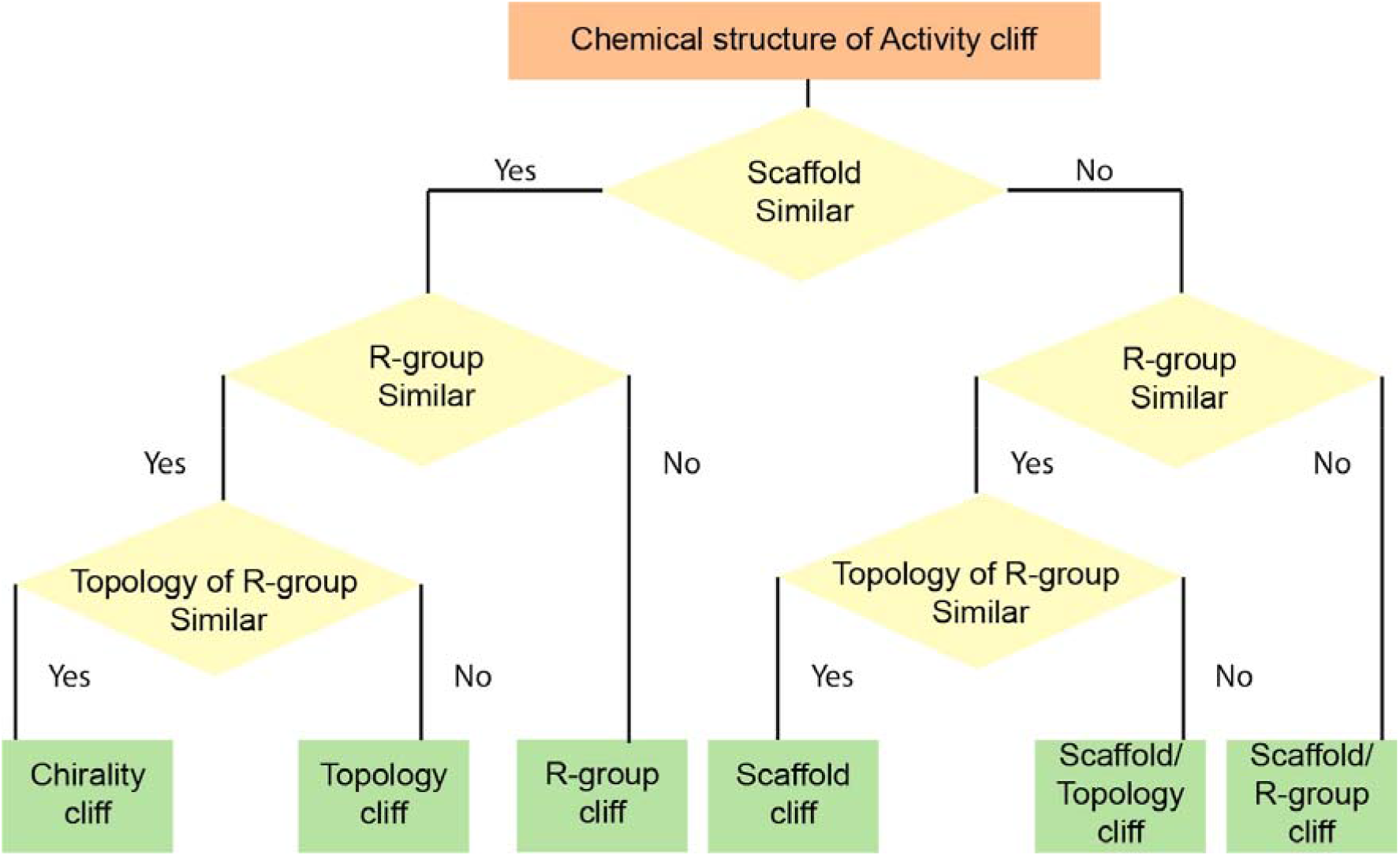
Computational workflow to structurally classify the activity cliff pairs using the structural information of the chemicals at different levels.

Note that the ‘Scaffold/R-group’ cliff category listed above was not considered by Hu et al. while defining the types of activity cliffs (Hu and Bajorath, 2012).

To classify the activity cliffs into six different structural categories, we used the R-group decomposition function in RDKit (“RDKit,” 2022), to decompose the chemical structure into its core structure (scaffold) and R-groups (Supplementary Table S4). Then using the workflow as depicted in Figure 7, we manually classified the 86 activity cliffs into 6 different structural categories (Supplementary Table S5). We find that among the 86 activity cliffs, 3 are Chirality cliffs, 3 are Topology cliffs, 29 are R-group cliffs, 2 are Scaffold cliffs, 9 are Scaffold/Topology cliffs, and 40 are Scaffold/R-group cliffs. In the next subsection, we illustrate with examples the six categories of activity cliffs by considering two ACGs and their activity cliff pairs.

### 3.7 Activity cliff classification of ACGs: DHT and 5α-Androstan-17β-ol

Here, we show the structural classification of the activity cliff pairs for two ACGs namely, DHT (CAS identifier: 521-18-6) and 5α-Androstan-17β-ol (CAS identifier: 1225-43-0). Figure 8 shows the activity cliff classification of the activity cliff pairs of the two ACGs. Further, Figure 8 also provides the CAS identifier and the logarithm of RBA value, i.e., log(RBA) of the chemicals which makeup the above activity cliff pairs.

**Figure 8:**
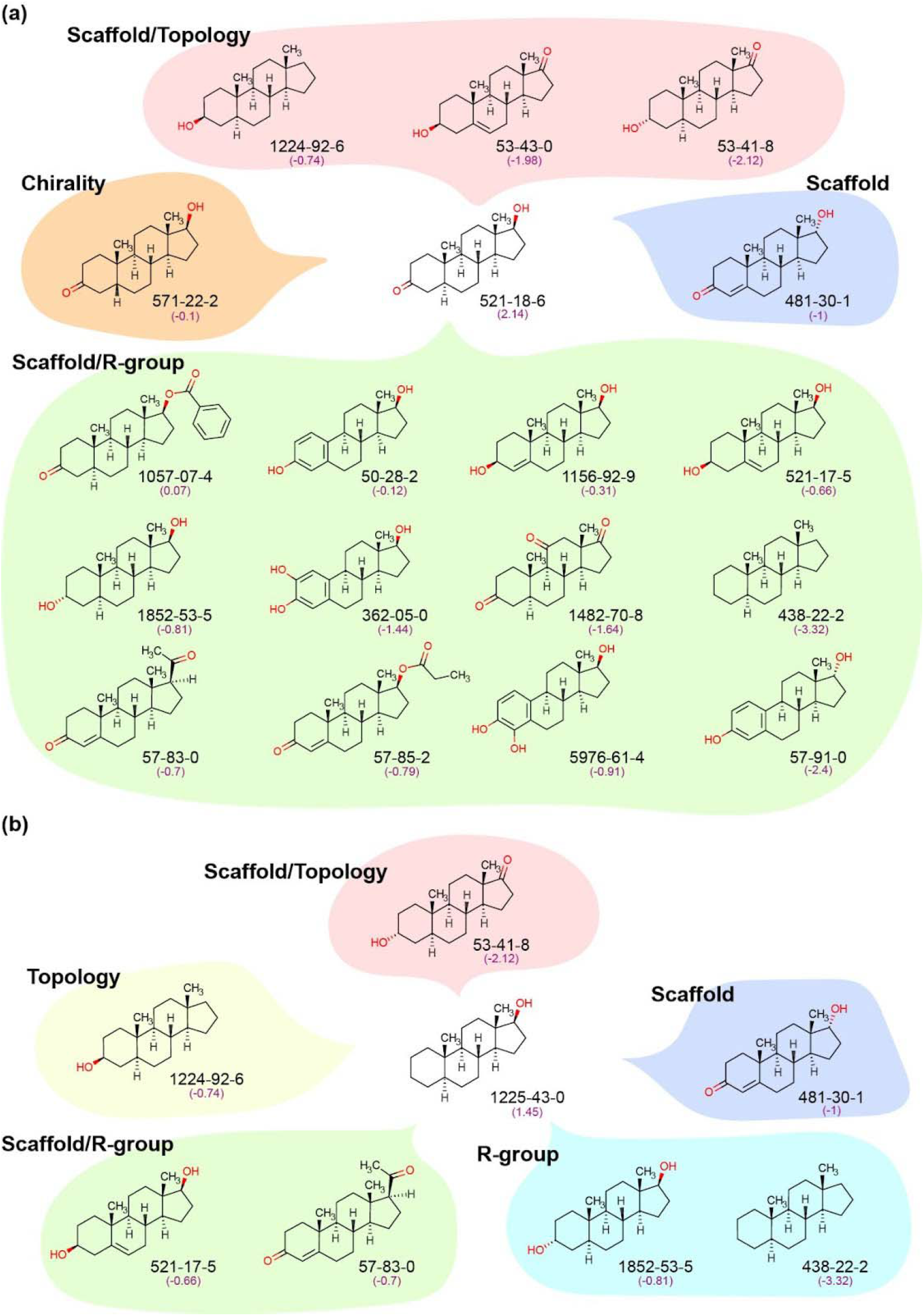
Structural classification of the Activity cliffs. **(a)** Structural classification of the 17 activity cliff pairs of the ACG DHT. **(b)** Structural classification of the 7 activity cliffs of the ACG 5α-Androstan-17β-ol. For each chemical, CAS identifier and log(RBA) values have been provided. Here, each activity cliff category is highlighted in a different color.

DHT is one of the principal androgens with high binding affinity to AR. DHT forms activity cliffs with 17 other chemicals, which is the maximum number of activity cliff pairs formed by any chemical in our dataset. All the 17 chemicals which form activity cliffs with DHT have low binding affinity compared to DHT. Using the computational workflow as shown in Figure 7, we were able to classify the 17 activity cliff pairs of DHT into 4 categories namely, Chirality cliff, Scaffold cliff, Scaffold/Topology cliff and Scaffold/R-group cliff (Figure 8a). Figure 8b shows the cliff classification of another ACG namely, 5α-Androstan-17β-ol forming activity cliffs with 7 other chemicals in our dataset. The 7 activity cliffs can be structurally classified into Topology cliff, R-group cliff, Scaffold cliff, Scaffold/Topology cliff and Scaffold/R-group cliff. Below, we explain the structural classification of a few activity cliff pairs for the two ACGs.

DHT and Etiocholan-17β-ol-3-one (CAS identifier: 571-22-2) are stereoisomers and form Chirality cliff, with activity difference (i.e., difference in log(RBA)) of 2.15 (Figure 8a). 5α-Androstan-17β-ol and 5α-Androstan-3β-ol (CAS identifier: 1224-92-6) form Topology cliff since both the chemicals have the same scaffold and R-groups (-CH_3_ and -OH), but only differ in the topology of the R-group (-OH), with activity difference of 2.19 (Figure 8b). 5α-Androstan-17β-ol forms R-group cliffs with two chemicals as these two chemicals when compared to 5α-Androstan-17β-ol have different R-groups but the same scaffolds (Figure 8b). 5α-Androstan-17β-ol forms Scaffold cliff with epitestosterone since both the chemicals differ in molecular scaffold but have the same R groups (-CH_3_, -OH) and topology of R-groups, with activity difference of 2.45 (Figure 8b). DHT forms Scaffold/Topology cliffs with three other chemicals which share the same set of R-groups (-CH3, -OH) with DHT, but have different molecular scaffolds and topology of R-groups (Figure 8a). DHT forms 12 Scaffold/R-group cliffs since these chemicals, when compared to DHT, differ in both molecular scaffolds and R-groups (Figure 8a).

## 4. Discussion and Conclusions

As part of our efforts to analyze the activity landscape of the AR binding chemicals, we employed several computational approaches to visualize and explore the chemical space, and to compare the global diversity of the whole library and its three identified clusters. Further, using SAS map based approach, we identified 86 activity cliffs in the whole library and found that the chemicals forming activity cliffs belong exclusively to one of the three clusters, i.e., cluster C2 which is dominated by ‘Steroids and steroid derivatives’. 14 chemicals that simultaneously form several activity cliffs were identified as ACGs. Additionally, a detailed inspection of the SALI heatmap revealed that most of the activity cliffs identified from the SAS map have high SALI scores. Lastly, we classified the activity cliffs into six categories by considering the chemical structure information of the AR binders at different levels. To the best of our knowledge, the present study is the first attempt to computationally analyze the structure-activity landscape of the AR binding chemicals.

In this study, we used three established computational approaches namely, a 2D visualization of the activity landscape (SAS map), a numerical scoring based approach (SALI), and structure based classification of activity cliffs, to expose the structure-activity landscape of AR binders (Guha and Van Drie, 2008; Hu and Bajorath, 2012; Medina-Franco et al., 2009; Méndez-Lucio et al., 2012; Naveja et al., 2018). While these three methods have individually been reported in published literature for the identification of activity cliffs, we here employed a combined approach leveraging the three methods to best characterize the structure-activity landscape of the AR binders. From our analysis, we observed that SAS map, which takes into account the pairwise comparison of structural similarity and activity difference, seems a better way to analyze the heterogeneity of the activity landscapes. Further, the SALI based approach was helpful in numerically quantifying the activity cliffs. Though some chemical pairs with high SALI scores were not identified as activity cliffs from the SAS map, the majority of the chemical pairs with high SALI scores were identified as activity cliffs. Further, we classified the identified activity cliffs into six categories based on the chemical structure information at different levels, and this facilitates better interpretation of the identified activity cliffs from a chemistry perspective (Hu and Bajorath, 2012).

In their analysis of the activity landscape of estrogen receptor binding chemicals, Naveja et al. had additionally interpreted some of the identified activity cliffs from SAS map using experimentally determined estrogen receptor protein structures co-crystallized with chemicals forming activity cliff pairs (Naveja et al., 2018). Naveja et al. had performed this analysis with the aim of elucidating the molecular mechanisms behind the activity difference between a pair of estrogen receptor binding chemicals constituting an activity cliff. Though we would have also liked to perform a similar analysis, unfortunately there were no available experimentally determined co-crystallized AR protein structures for rat or mouse or human in the Protein Data Bank (PDB; https://www.rcsb.org/) for any pair of AR binding chemicals that constitute an activity cliff identified from SAS map. On the other hand, we present the structural categorization of the activity cliffs proposed by Hu and Bajorath to aid in structural interpretation of the activity cliffs (Hu and Bajorath, 2012).

In conclusion, we present a comprehensive analysis of the structure-activity landscape of a benchmark dataset of 144 AR binding chemicals by leveraging multiple complementary computational approaches. To reiterate, this is the first study to perform detailed activity landscape analysis of the AR binding chemicals. We expect that the insights from this study will help chemists in better interpretation of the structural features behind activity cliffs, and moreover, will facilitate ongoing attempts in the area of computational toxicology towards building predictive models for identifying endocrine or hormone disruptors in the ever-expanding human chemical exposome.

## Acknowledgements

Areejit Samal acknowledges support from the Department of Atomic Energy, Government of India and a Max Planck Partner Group in Mathematical Biology funded by the Max Planck Society Germany. The funders have no role in study design, data collection, data analysis, manuscript preparation or decision to publish.

## CRediT author contribution statement

**R.P. Vivek-Ananth:** Conceptualization, Data curation, Software, Formal analysis, Visualization, Writing. **Ajaya Kumar Sahoo:** Conceptualization, Data curation, Software, Formal analysis, Visualization, Writing. **Shanmuga Priya Baskaran:** Conceptualization, Data curation, Software, Formal analysis, Visualization, Writing. **Janani Ravichandran:** Formal analysis, Writing. **Areejit Samal:** Conceptualization, Supervision, Formal analysis, Writing.

## Supplementary Tables

**Table S1**: Curated list of 144 AR binding chemicals analyzed in this study. For each chemical, the CAS identifier and binding affinity in terms of log(RBA) were collected from Fang et al. The chemical structures are presented in Canonical SMILES notation. Further, for each chemical, we provide the Bemis-Murcko scaffolds computed using RDKit, chemical class information predicted from ClassyFire, and chemical cluster information from CSN.

**Table S2**: Computed SALI scores for 86 activity cliff pairs identified from the SAS map. The chemicals are identified by their CAS identifiers. The pairwise structural similarity for each activity cliff pair is computed via Tanimoto coefficient using ECFP4 fingerprints. The activity difference for each activity cliff pair is computed as the absolute difference of the log(RBA) values. Note that, the Tanimoto similarity value is shown here up to 2 decimal places.

**Table S3**: List of 14 activity cliff generators (ACGs) identified from the SAS map. The ACGs are identified by their CAS identifiers. For each ACG, we provide the chemicals (CAS identifiers) separated by ‘|’ symbol with which the ACG forms activity cliffs. Further, we provide the number of activity cliff pairs formed by each ACG.

**Table S4**: R-group decomposition of the 41 chemicals identified to form 86 activity cliff pairs in the SAS map. The chemicals are identified by their CAS identifiers. For each chemical, we show the core structure (scaffold) and the R-groups (separated by ‘|’ symbol) identified using the R-group decomposition function in RDKit.

**Table S5**: Structural classification of the 86 activity cliff pairs into six categories using the computational workflow shown in Figure 7 that takes into account the structural information of the chemicals at different levels. The chemicals are identified by their CAS identifiers.

## References

Bajorath, J., 2017. Representation and identification of activity cliffs. Expert Opinion on Drug Discovery 12, 879–883. https://doi.org/10.1080/17460441.2017.1353494

Bastian, M., Heymann, S., Jacomy, M., 2009. Gephi: An Open Source Software for Exploring and Manipulating Networks. ICWSM 3, 361–362. https://doi.org/10.1609/icwsm.v3i1.13937

Bemis, G.W., Murcko, M.A., 1996. The Properties of Known Drugs. 1. Molecular Frameworks. J. Med. Chem. 39, 2887–2893. https://doi.org/10.1021/jm9602928

Blondel, V.D., Guillaume, J.-L., Lambiotte, R., Lefebvre, E., 2008. Fast unfolding of communities in large networks. Journal of Statistical Mechanics: Theory and Experiment 2008, P10008. https://doi.org/10.1088/1742-5468/2008/10/P10008

Cruz-Monteagudo, M., Medina-Franco, J.L., Pérez-Castillo, Y., Nicolotti, O., Cordeiro, M.N.D.S., Borges, F., 2014. Activity cliffs in drug discovery: Dr Jekyll or Mr Hyde? Drug Discovery Today 19, 1069–1080. https://doi.org/10.1016/j.drudis.2014.02.003

Davey, R.A., Grossmann, M., 2016. Androgen Receptor Structure, Function and Biology: From Bench to Bedside. Clin Biochem Rev 37, 3–15.

Djoumbou Feunang, Y., Eisner, R., Knox, C., Chepelev, L., Hastings, J., Owen, G., Fahy, E., Steinbeck, C., Subramanian, S., Bolton, E., Greiner, R., Wishart, D.S., 2016. ClassyFire: automated chemical classification with a comprehensive, computable taxonomy. Journal of Cheminformatics 8, 61. https://doi.org/10.1186/s13321-016-0174-y

Fang, H., Tong, W., Branham, W.S., Moland, C.L., Dial, S.L., Hong, H., Xie, Q., Perkins, R., Owens, W., Sheehan, D.M., 2003. Study of 202 Natural, Synthetic, and Environmental Chemicals for Binding to the Androgen Receptor. Chem. Res. Toxicol. 16, 1338–1358. https://doi.org/10.1021/tx030011g

González-Medina, M., Owen, J.R., El-Elimat, T., Pearce, C.J., Oberlies, N.H., Figueroa, M., Medina-Franco, J.L., 2017. Scaffold Diversity of Fungal Metabolites. Frontiers in Pharmacology 8, 180. https://doi.org/10.3389/fphar.2017.00180

González-Medina, M., Prieto-Martínez, F.D., Owen, J.R., Medina-Franco, J.L., 2016. Consensus Diversity Plots: a global diversity analysis of chemical libraries. Journal of Cheminformatics 8, 63. https://doi.org/10.1186/s13321-016-0176-9

Guha, R., 2012. Exploring structure–activity data using the landscape paradigm. WIREs Computational Molecular Science 2, 829–841. https://doi.org/10.1002/wcms.1087

Guha, R., Van Drie, J.H., 2008. Structure-Activity Landscape Index:□ Identifying and Quantifying Activity Cliffs. J. Chem. Inf. Model. 48, 646–658. https://doi.org/10.1021/ci7004093

Hu, Y., Bajorath, J., 2012. Extending the Activity Cliff Concept: Structural Categorization of Activity Cliffs and Systematic Identification of Different Types of Cliffs in the ChEMBL Database. J. Chem. Inf. Model. 52, 1806–1811. https://doi.org/10.1021/ci300274c

Hunter, J.D., 2007. Matplotlib: A 2D Graphics Environment. Computing in Science & Engineering 9, 90–95. https://doi.org/10.1109/MCSE.2007.55

Jeng, H.A., 2014. Exposure to Endocrine Disrupting Chemicals and Male Reproductive Health. Frontiers in Public Health 2, 55. https://doi.org/10.3389/fpubh.2014.00055

Jolliffe, I.T., 1986. Principal Component Analysis, Springer Series in Statistics. Springer New York, New York, NY. https://doi.org/10.1007/978-1-4757-1904-8

Karthikeyan, B.S., Ravichandran, J., Aparna, S.R., Samal, A., 2021. DEDuCT 2.0: An updated knowledgebase and an exploration of the current regulations and guidelines from the perspective of endocrine disrupting chemicals. Chemosphere 267, 128898. https://doi.org/10.1016/j.chemosphere.2020.128898

Karthikeyan, B.S., Ravichandran, J., Mohanraj, K., Vivek-Ananth, R.P., Samal, A., 2019. A curated knowledgebase on endocrine disrupting chemicals and their biological systems-level perturbations. Science of The Total Environment 692, 281–296. https://doi.org/10.1016/j.scitotenv.2019.07.225

Lipkus, A.H., Yuan, Q., Lucas, K.A., Funk, S.A., Bartelt, W.F.I., Schenck, R.J., Trippe, A.J., 2008. Structural Diversity of Organic Chemistry. A Scaffold Analysis of the CAS Registry. J. Org. Chem. 73, 4443–4451. https://doi.org/10.1021/jo8001276

Maggiora, G.M., 2006. On Outliers and Activity CliffsWhy QSAR Often Disappoints. J. Chem. Inf. Model. 46, 1535. https://doi.org/10.1021/ci060117s

Medina-Franco, J.L., Martínez-Mayorga, K., Bender, A., Marín, R.M., Giulianotti, M.A., Pinilla, C., Houghten, R.A., 2009. Characterization of Activity Landscapes Using 2D and 3D Similarity Methods: Consensus Activity Cliffs. J. Chem. Inf. Model. 49, 477–491. https://doi.org/10.1021/ci800379q

Méndez-Lucio, O., Pérez-Villanueva, J., Castillo, R., Medina-Franco, J.L., 2012. Identifying Activity Cliff Generators of PPAR Ligands Using SAS Maps. Molecular Informatics 31, 837–846. https://doi.org/10.1002/minf.201200078

Morgan, H.L., 1965. The Generation of a Unique Machine Description for Chemical Structures-A Technique Developed at Chemical Abstracts Service. J. Chem. Doc. 5, 107–113. https://doi.org/10.1021/c160017a018

Naveja, J.J., Medina-Franco, J.L., 2015. Activity landscape sweeping: insights into the mechanism of inhibition and optimization of DNMT1 inhibitors. RSC Adv. 5, 63882–63895. https://doi.org/10.1039/C5RA12339A

Naveja, J.J., Norinder, U., Mucs, D., López-López, E., Medina-Franco, J.L., 2018. Chemical space, diversity and activity landscape analysis of estrogen receptor binders. RSC Adv. 8, 38229–38237. https://doi.org/10.1039/C8RA07604A

Peltason, L., Bajorath, J., 2007. SAR Index:□ Quantifying the Nature of Structure-Activity Relationships. J. Med. Chem. 50, 5571–5578. https://doi.org/10.1021/jm0705713 RDKit, 2022.

Rehman, Saba, Usman, Z., Rehman, Sabeen, AlDraihem, M., Rehman, N., Rehman, I., Ahmad, G., 2018. Endocrine disrupting chemicals and impact on male reproductive health. Translational Andrology and Urology 7, 490–503. https://doi.org/10.21037/tau.2018.05.17

Rodprasert, W., Toppari, J., Virtanen, H.E., 2021. Endocrine Disrupting Chemicals and Reproductive Health in Boys and Men. Frontiers in Endocrinology 12, 706532. https://doi.org/10.3389/fendo.2021.706532

Rogers, D., Hahn, M., 2010. Extended-Connectivity Fingerprints. J. Chem. Inf. Model. 50, 742–754. https://doi.org/10.1021/ci100050t

Shanmugasundaram, V., Maggiora, G., 2001. Characterizing property and activity landscapes using an information-theoretic approach. 222nd American Chemical Society National Meeting, Division of Chemical Information, p. Abstract No. 77.

Sud, M., 2016. MayaChemTools: An Open Source Package for Computational Drug Discovery. J. Chem. Inf. Model. 56, 2292–2297. https://doi.org/10.1021/acs.jcim.6b00505

Tan, M.E., Li, J., Xu, H.E., Melcher, K., Yong, E., 2015. Androgen receptor: structure, role in prostate cancer and drug discovery. Acta Pharmacologica Sinica 36, 3–23. https://doi.org/10.1038/aps.2014.18

Tanimoto, T. T., 1957. IBM Internal Report 17th Nov.

UNEP, W., 2013. State of the science of endocrine disrupting chemicals-2012. WHO-UNEP, Geneva.

Vivek-Ananth, R.P., Sahoo, A.K., Baskaran, S.P., Samal, A., 2022. Scaffold and structural diversity of the secondary metabolite space of medicinal fungi. bioRxiv 2022.09.25.509364. https://doi.org/10.1101/2022.09.25.509364

Wassermann, A.M., Wawer, M., Bajorath, J., 2010. Activity Landscape Representations for Structure-Activity Relationship Analysis. J. Med. Chem. 53, 8209–8223. https://doi.org/10.1021/jm100933w

Wawer, M., Peltason, L., Weskamp, N., Teckentrup, A., Bajorath, J., 2008. Structure-Activity Relationship Anatomy by Network-like Similarity Graphs and Local Structure-Activity Relationship Indices. J. Med. Chem. 51, 6075–6084. https://doi.org/10.1021/jm800867g

